# Self-Organized Pattern Formation of a Common Tropical Alga

**DOI:** 10.64898/2026.01.13.699319

**Authors:** Dylan McNamara, Conner Lester, Clinton Edwards, Jennifer Smith, Stuart Sandin

## Abstract

Recent in-situ observations within a tropical coral reef have revealed novel polygonal patterns of the calcifying green alga *Halimeda*. The observed patterns showed no evidence of a matching exogenous template in structural reef morphology, distribution of biological competitors for space, or other environmental factors, suggesting the pattern is the result of endogenous, nonlinear pattern forming dynamics. A simplified, spatially explicit numerical model is proposed that simulates a feedback whereby *Halimeda* preferentially grows in regions less conducive to the growth of corals (when corals are the dominant spatial competitor), and coral growth is inhibited in regions of dense *Halimeda*. Model results reveal self-organized emergent polygons of *Halimeda* cover that qualitatively match observations.

## 1 Introduction

Striking, regular patterns have been observed in a wide range of ecological environments. Examples can be found in arid, wetland, savannah, mudflat, marine, and forest ecosystems ([1],[2],[3]). The most common explanation for the dynamics of pattern formation is a process known as scale-dependent feedback (SDF) ([4]), also initially referred to as Turing instabilities ([5]), whereby short-range enhancement and long-range depletion of a system characteristic leads to the self-organized emergence of a coherent pattern on space and time scales larger than those that govern the local dynamics.

However, in some cases of observed ecosystem patterns, scale-dependent feedbacks are not found, and an alternative explanation has been suggested. This mechanism is termed density-dependent aggregation (DDA) of an abiotic or biotic feature and can give rise to large-scale patches in space ([6]). These patches compete for resources, and this dynamical competition results in a slow time-scale coarsening of patch size as larger patches outcompete smaller ones. In contrast to SDF, the resulting patterns from DDA are not regularly spaced and do not reach steady state until a single winning patch remains.

Notably, the preceding examples may not provide the only mechanisms leading to self-organized and emergent patterns within biological systems. Recent observations using reconstructed images of underwater coral reefs revealed a novel pattern in the common tropical alga *Halimeda* (Figure 1 [7]). The patterns in this alga are distinct from the typical patterns resulting from SDF and DDA. Namely, the *Halimeda* are not clumped or striped with a fixed wavelength (SDF) and they are not in patches of various sizes (DDA). Instead they were found to be organized into polygons with a somewhat consistent wavelength.

**Fig. 1.**
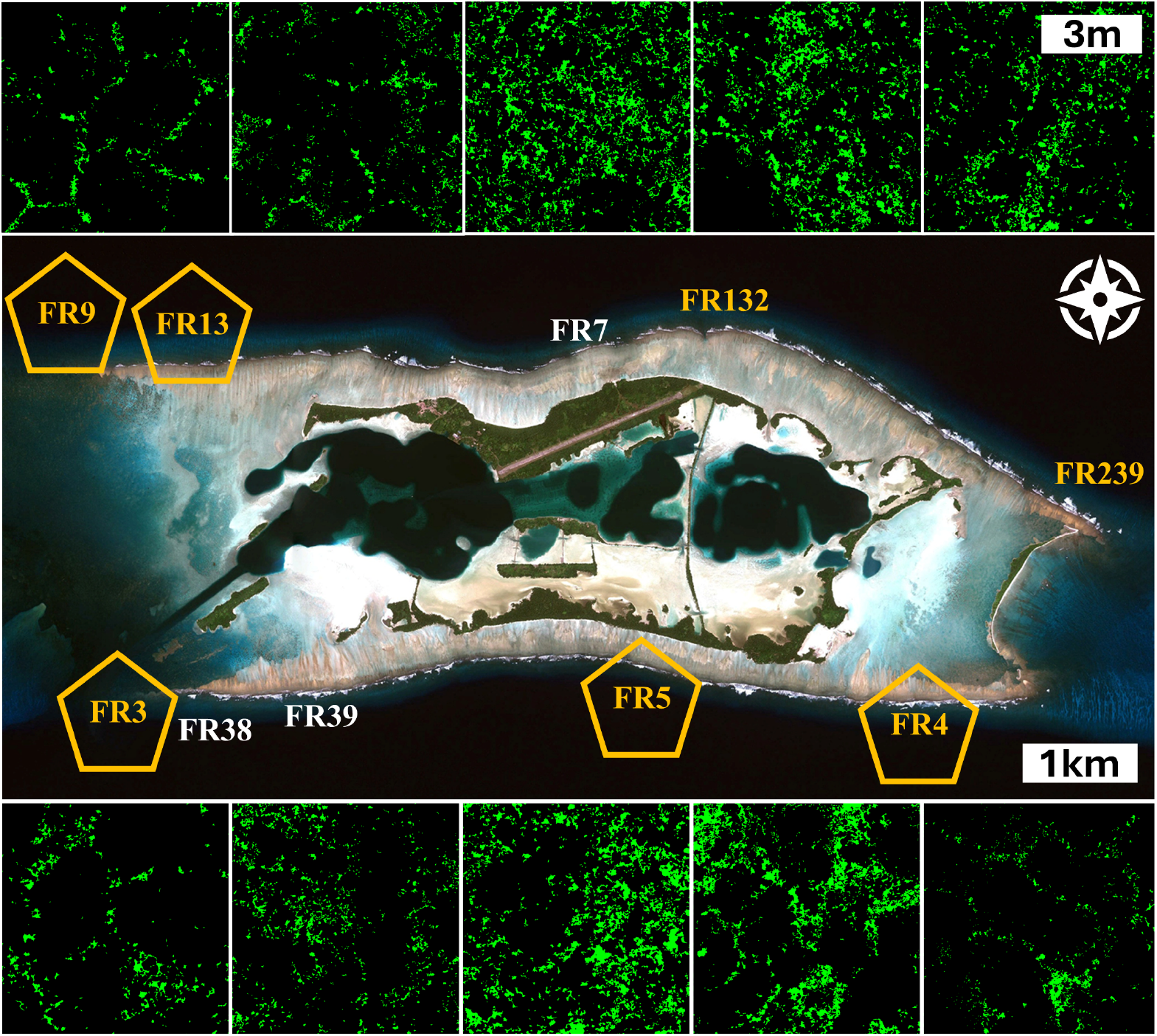
Polygonal patterns of *Halimeda* at five out of ten indicated locations around Palmyra atoll in the Northern Line Islands. Each image is 100 *m*^2^ and has been generated by merging approximately 5000 overlapping photographs of the coral benthos into a single orthomosaic. The resulting image is then annotated to identify individual pixels as either *Halimeda* (green) or not (black). See Sandin et al. [7] for more information about the orthomosaic process.

Building from pioneering work on pattern formation in geomorphology where self-organization was shown to give rise to a variety of patterns in sorted ground ([8]), here we propose a numerical model to explain how *Halimeda* polygons can form in coral reefs. The feedbacks that drive self-organization in our model occur between characteristics of dominant spatial competitors (in this case, stony corals) and *Halimeda*. The general form of such feedbacks has been recently coined scale dependent feedbacks in time (SDFt) ([9]) to contrast with scale dependent feedbacks which occur in space (SDFs), like the aforementioned Turing instabilities.

Investigating how patterns form in natural systems provides insight into endogenous system processes and when that foundation is strengthened, there is renewed understanding in how the given system will respond to exogenous disturbance. In the case of coral reefs, which are increasingly stressed due to both local and global human impacts, breakthroughs in understanding the feedbacks between coral and algae that cause pattern formation increase insight into stability of coral reefs and hence can help inform future conservation efforts.

## 2 Model

The model evolves two variables in space and time; *Halimeda* biomass (H) and the quality of the benthos for recruitment and growth of coral (Q), which we will refer to simply as coral quality. The former variable is a well-defined measurable quantity. The latter is a proxy that is meant to capture aspects of the benthos such as structural consolidation ([10]) and health of the microbial community([11]).

Temporal changes in *Halimeda* biomass are represented by three processes described as follows:

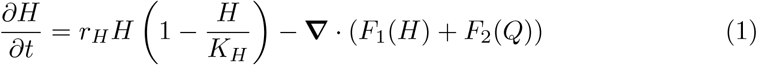

The first term on the right simulates logistic growth, at rate *r*_*H*_, to a carrying capacity, *K*_*H*_. This is a common abstraction of biological growth processes and captures the growth of *Halimeda* at a given location in the absence of competitors. The next term shows the divergence of two types of *Halimeda* fluxes, one that depends on the biomass of *Halimeda, F*_1_(*H*), and another that is related to coral quality, *F*_2_(*Q*). These two terms are representations of the spreading of *Halimeda* across space.

The flux term *F*_1_(*H*) simulates spatial movement of *Halimeda* due to the combined impact of negative and positive growth ([12]). When pieces of *Halimeda* fragment and die, they then become a focal point for new growth, and in this sense, *Halimeda* spread in the direction that pieces fall. As noted, this flux depends on the local amount of *Halimeda*, so we represent that flux as

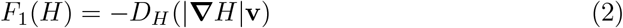

where *D*_*H*_ is a parameter controlling the strength of *Halimeda* flux. The term **v** is a unitless vector representing the direction of *Halimeda* flux, or the favored direction that *Halimeda* spread in space. Our model explores two choices for this vector; either a unit vector pointing along the axis of surrounding *Halimeda* (where the axis is defined over a distance *L* centered on the given location) or a unit vector oriented along the gradient in *Halimeda* biomass. With the former choice, the spatial spread of *Halimeda* mimics linear rhizoidal growth ([13]). The latter choice leads to the spreading of *Halimeda* as would happen from a typical isotropic diffusive process.

The term *F*_2_(*Q*) is related to coral quality and moves *Halimeda* away from regions where the benthos is instead more suitable for coral growth. This is accomplished by letting

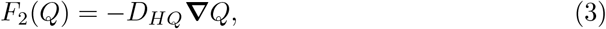

where *D*_*HQ*_ is a diffusion constant. This term has abstracted a number of complex processes related to microbially mediated competition between algae and corals. The net impact of these processes has been empirically determined to separate regions of coral from regions of *Halimeda* ([14]). The mathematical representation simulates this observed dynamic as *Halimeda* will flux away from regions of high coral quality (and the associated microbial community) to regions with lower coral quality. Substituting these fluxes into the *Halimeda* evolution equation gives,

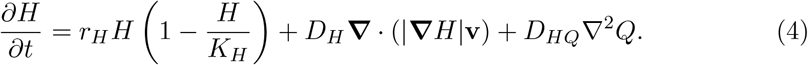

Temporal changes in coral quality arise from two processes in our model. Coral quality grows at a given spatial location to a maximum value that is inversely related to the amount of surrounding *Halimeda*, so that more *Halimeda* results in a smaller maximum allowable coral quality. Coral quality also diffuses spatially to simulate the random processes that distribute aspects of benthic coral quality such as the spread of crustose coraline algae or microbial communities that are conducive to coral growth. These processes are represented as

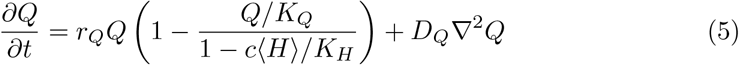

where *r*_*Q*_ is the growth rate of coral quality, *K*_*Q*_ is coral quality capacity, *c* is a parameter that scales the size of the maximum coral quality, and *D*_*Q*_ is a diffusion constant. The maximum allowable coral quality is found by averaging *Halimeda* over a neighborhood as

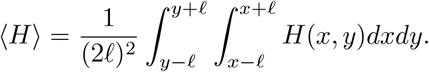

The equations above contain two terms that couple *Halimeda* and coral quality and provide an avenue for feedbacks that can cause a pattern to self-organize. The term *D*_*HQ*_∇ ^2^*Q* in Eq. 4 causes *Halimeda* to collect in regions removed from high coral quality and in concert with the first term of Eq. 5, these zones of *Halimeda* aggregation lead to a decrease in the maximum allowable coral quality. As coral quality decreases where *Halimeda* biomass is large, it leads to stronger gradients in coral quality that continue to drive *Halimeda* to collect in increasingly dense regions. Overall this leads to *Halimeda* attracting *Halimeda* and coral quality in the benthos promoting higher coral quality, with both separating from each other. A second potential feedback occurs via the second term in Eq. 4 when **v** is chosen to orient along the long axis of surrounding *Halimeda*. This dynamic squeezes *Halimeda* along an axis, thereby promoting increases in surrounding coral quality that elongate *Halimeda* axes even further. In total, these two feedbacks and the other dynamics are novel partial differential equation representations of similar processes operating in permafrost regions ([8]).

The model equations above abstract detailed processes occurring both in *Halimeda* growth and in the benthic environment as it relates to the potential for coral occupation and growth. For example, *Halimeda* is simply assumed to grow to a carrying capacity with a preference for elongating when **v** is chosen appropriately ([13]). The details of the interaction between coral quality and *Halimeda* have been collectively represented as spatially separating the two from each other ([14]). The goal of all of these abstractions is to reduce the dynamics of and between coral quality and *Halimeda* to their simplest essence and then to explore whether such dynamics can cause patterns that are similar to those found in nature ([7]). Further, while the observed polygonal patterns in *Halimeda* are clear, the benthos within the observed polygons is less easily characterized. There are no consistent species or elevations within the polygons ([7]). In other words, when looking at taxonomically classified images inside a region bordered by a *Halimeda* polygon, there is a distribution of varying species rather than just one. And the elevation of the benthos inside *Halimeda* polygons is not fixed within each nor across many. This absence of an obvious environmental template suggests the *Halimeda* have self-organized.

As the postulated dynamics in Eqs. 4 and 5 are simplified abstractions, some of the parameters cannot be easily grounded empirically. To reduce the number of independently varying parameters that might impact pattern formation, Eqs. 4 and 5 are nondimensionalized using the following scalings,

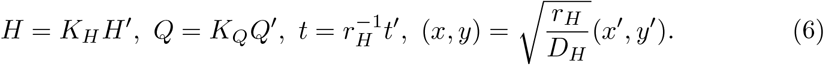

Scaling all of the variables reduces the number of free coefficient parameters to five (with a sometimes sixth parameter in *L*). The resulting nondimensional equations are

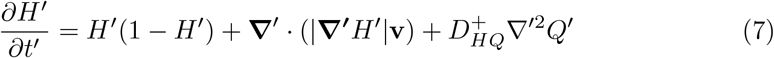

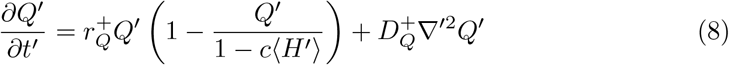

where 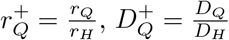, and 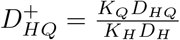.

As mentioned, setting **v** so that the velocity term is oriented in the direction of the gradient in *Halimeda*, reduces the second term on the right hand side of Eq. 7 to linear diffusion ∇^*′*2^*H*^*′*^ of *Halimeda*. This simplification allows a straightforward linear stability analysis of the systems of equations in Eqs. 7 and 8. Details of this analysis are left to the appendix.

## 3 Results

The linear stability analysis confirms that a spatially homogeneous equilibrium is unstable for certain parameter combinations (Fig. A1). To confirm the instability and explore the resulting pattern that forms at finite amplitude, we numerically solve the governing equations 7 and 8 starting from random perturbations on the steady state homogeneous equilibrium solution *H*^*′*^ = 1, *Q*^*′*^ = 1 − *c*. The domain in this and all other simulations is periodic. Unless otherwise noted, *l* = 0.75 as size of the area for averaging *Halimeda* and *L* = 2.5 for defining the *Halimeda* axis. As a starting point, we establish a canonical setting for the model parameters of 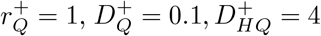, and *c* = 0.6. Figure 2 shows the evolution of both *H*^*′*^ and *Q*^*′*^ from the initial state to the steady state pattern for the case of **v** being oriented along the gradient in *Halimeda*, which again is simply spreading *Halimeda* in an isotropic diffusive process.

**Fig. 2.**
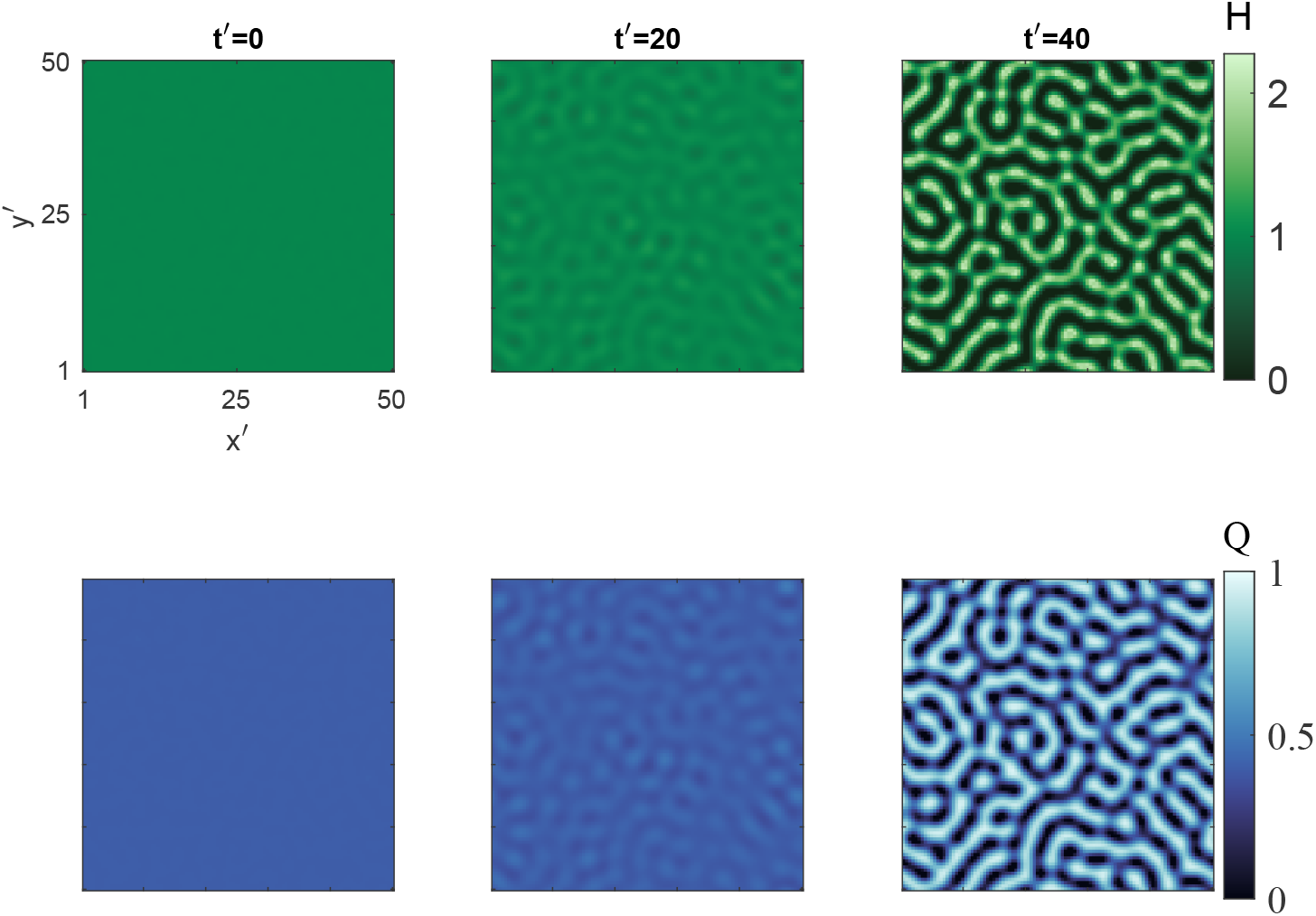
Snapshots from the model domain showing the evolution *Halimeda*, H (top), and coral quality, Q (bottom). From left to right the snapshots occur at t’=1,20,40

Setting **v** so that it is oriented along the long axis of surrounding *Halimeda* results in a steady state pattern in which the regions of *Halimeda* are squeezed tighter into polygons (Figure 3). The wavelength of the resulting pattern in this case is slightly smaller than when *Halimeda* spreads isotropically. This is because diffusing *Halimeda* preferentially along its surrounding long axis has a dynamically similar effect to increasing 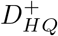 in Eq. 7. And as noted in the linear stability analysis (Fig. A1), increasing 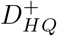 reduces the most unstable wavelength. This effective change to 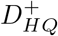 also increases the growth rate of the instability and indeed the polygonal pattern in this case forms faster than the patterns in the isotropic diffusion case (Figure 2).

**Fig. 3.**
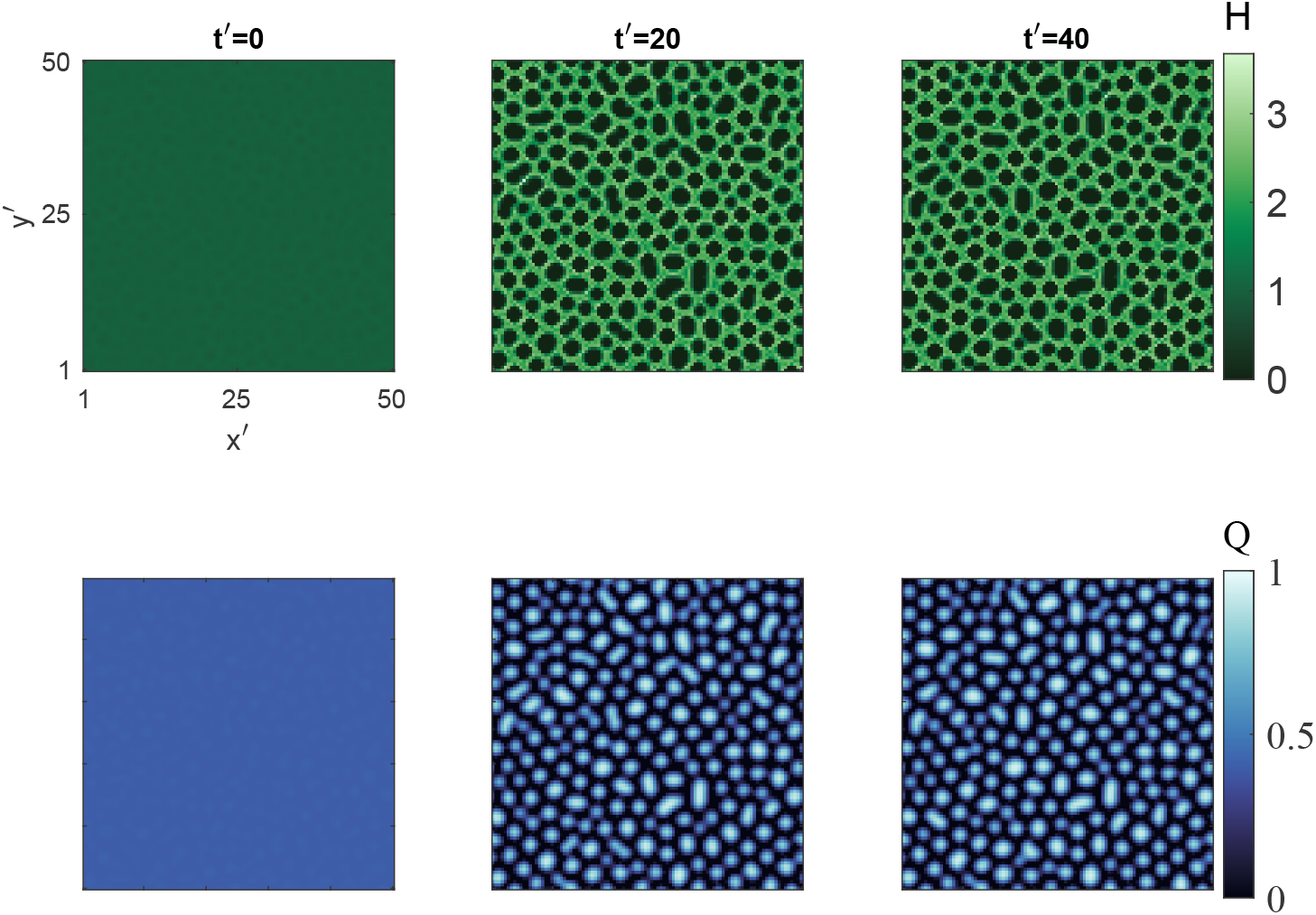
Snapshots from the model domain showing the evolution *Halimeda*, H (top), and coral quality, Q (bottom). From left to right the snapshots occur at t’=1,20,40. In this case **v** has been oriented along the local *Halimeda* axis and *L*=2.5.

The expected variation (from the linear stability analysis) in the resulting pattern characteristics as *c* and 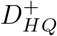 increases is matched by the model in finite amplitude (see the Supplementary section for further exploration of the pattern behavior over the model parameter space). Figure 4 shows that when these parameters are small no pattern forms but once they cross a threshold value, a pattern forms. The polygonal styled pattern is evident only when **v** is oriented along the long axis of surrounding *Halimeda*. As both *c* and 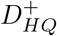 are increased further, the wavelength of the resulting pattern is reduced. These two parameters control the strength of the coupling between coral quality and *Halimeda*. In the field (Figure 1), polygons are more commonly found in regions with milder wave conditions ([7]). When waves are more frequently large, the spatially random wave disturbances would likely reduce the coupling strength between coral quality and *Halimeda* and according to our model results, no pattern would be expected to form (Figure 4). Furthermore, a competitive advantage is conferred to *Halimeda* via an increase in *K*_*H*_, which leads to a decrease in 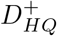 and a region of parameter space with no pattern formation. This could result for example, from increased nutrient levels that favor *Halimeda* and indeed the regions with no pattern in Figure 1 have considerably more *Halimeda*.

**Fig. 4.**
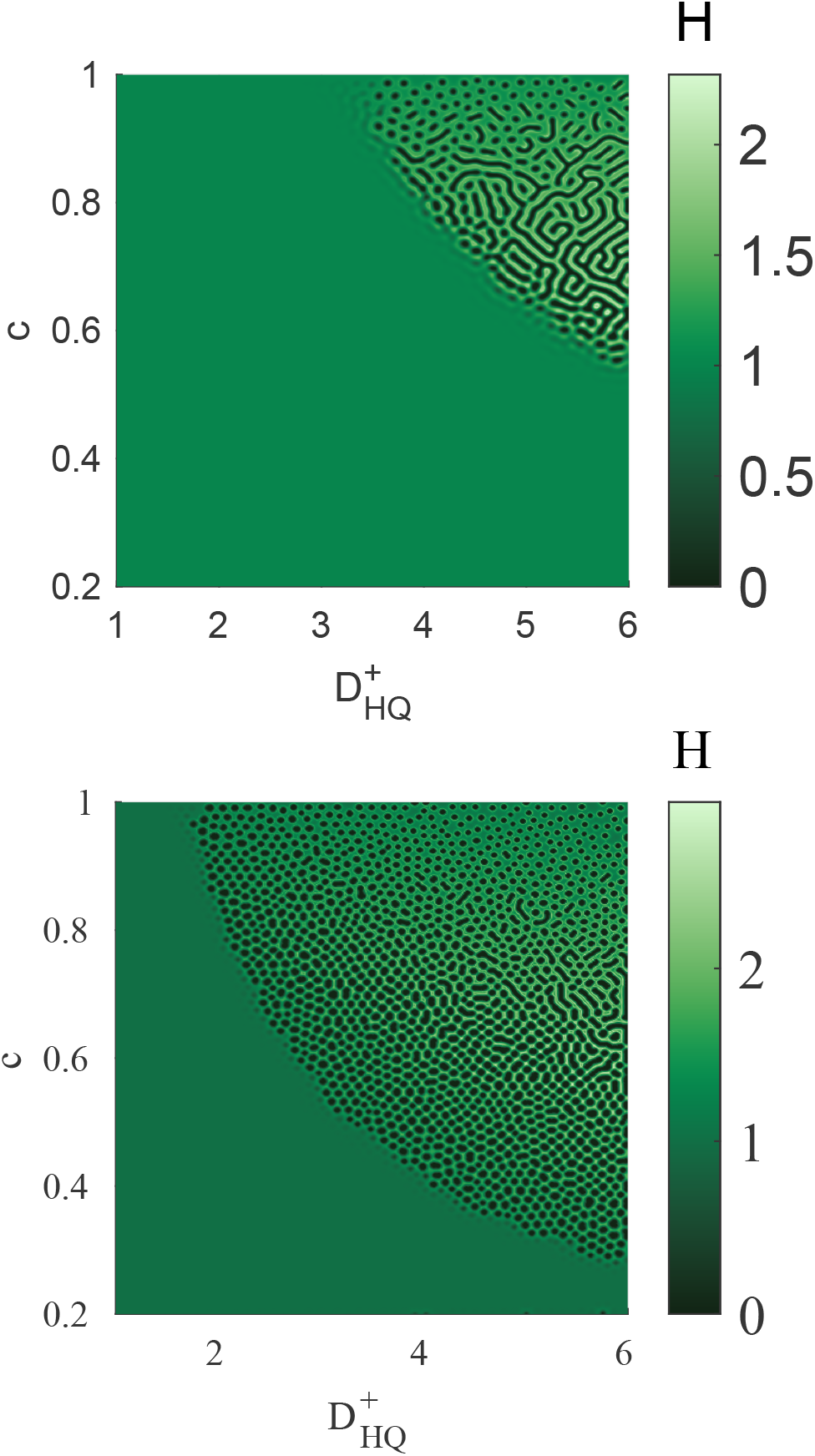
Model simulations that show variation in the steady state pattern’s existence and wavelength with changes to the nondimensional parameters *c* and 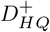 for when the velocity **v** is oriented along the gradient in *Halimeda* (top) and when it is oriented along the axis of surrounding *Halimeda* (bottom). For these simulations the size of the area, *l*, for averaging *Halimeda* is set to 1.25.

## 4 Conclusion

The presented model abstracts some of the detailed processes associated with *Halimeda* growth and the dynamical evolution of biologically created patterns in coral reef benthic environments. The model equations also include parameters whose values are unconstrained. The abstractions help to minimize postulated model dynamics, which in this case is particularly advantageous due to the paucity of measurements at time and space scales of the observed patterns in *Halimeda*. The simplifications of detailed processes also has the benefit of keeping the list of unknown parameters to a minimum.

Despite these necessary caveats, our simple model qualitatively reproduces the observed patterns in *Halimeda*. The steady-state model pattern when *Halimeda* grows along its surrounding axis (Figure 3) is similar in shape and scale to the observed patterns (Figure 1). Additionally, the model matches the observation that *Halimeda* polygons are found in regions of low wave conditions which would be more conducive to strong spatial coupling between coral quality and *Halimeda*. The modeled pattern also agrees with observations in the field that show the polygonal patterns remain intact over yearly time scales ([7]). Imposed perturbations to the modeled pattern do not alter the pattern location (not shown) unless perturbations are so significant that the pattern is effectively removed. This spatial memory is related to the long-time scale emergent dynamics of the pattern, whereby the dynamics of the pattern itself are scale separated from the faster-scale dynamics of the individual processes. Field observations suggest that the emergent pattern time-scale is at least many years to a decade as polygonal features have remained intact over that span. Maintaining the ongoing observational record is critical to refining this measure.

The model presented here can be easily generalized to other systems with polygonal features. This was similarly true in the early pioneering work on sorted patterned ground ([8]), but as that model was rule-based with rules that act on discrete entities (stones), it did not lend itself to systems with continuous scalar fields. In addition, our model framework is amenable to relatively simple analytical exploration of global trends in initial instability and pattern wavelength.

Recent authors have proposed the idea that spatial dynamics associated with pattern formation hold potential for increasing system resilience ([15]) via the muting of catastrophic phase shifts. For coral reefs, which are subject to increasing external stress and have been postulated to catastrophically shift to being dominated by macroalgae, this means that insight into pattern formation may inform reef resilience.

This is particularly true in the case of *Halimeda*, which is a natural component of coral reef systems and is a large contributor to carbonate production. With complementary observations of spatial patterns in benthic organisms, the stylized model presented here is a first step towards an increased understanding of the types of spatial dynamics that may lead to enhanced resilience of coral reefs.

## Acknowledgements

We thank the Nature Conservancy for logistical support and the US Fish and Wildlife Service for special use permit 12533-13025 and access to the refuge. This work is a contribution of the Reefs Tomorrow Initiative, a program funded by the Gordon and Betty Moore Foundation (grant 3420).

## Declarations

## Author contribution

All authors conceived of the study idea. DEM developed the mathematical model and along with CWL carried out the analysis. DEM wrote the first draft with input from SAS. All authors contributed to revisions and the final manuscript.

## Appendix A Linear Stability Analysis

Dropping the prime notation from Eqs. 7 and 8 and assuming that the velocity **v** is randomly oriented such that the second term in Eq. 7 reduces to diffusion, yields the following system of equations,

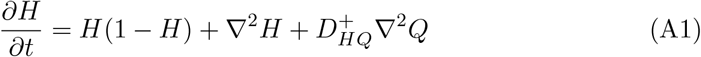

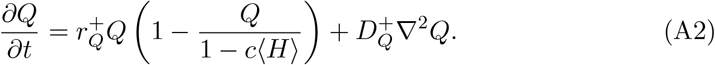

We perform a linear stability analysis about the fixed points (*H*^*^, *Q*^*^) = (1, 1 − *c*).

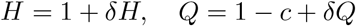

Plugging these expressionis into A1 and A2, then keeping everything to first order we find

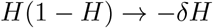

and

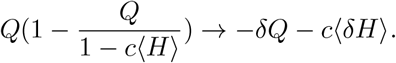

We explore Fourier modes of the perturbations *δH, δQ* ∼ *e*^*σt*^*e*^*ikx*^*e*^*iky*^ where *σ* is the growth rate (rescaled by *r*_*H*_) and *k* = 2*π/λ* is the wavenumber (rescaled by 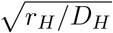).

These definitions give

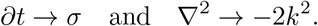

The spatial average of *δH* is now

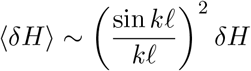

which reduces to ⟨*δH*⟩ ∼ *δH* as *k*ℓ → 0.

We now have the eigenvalue problem L***v*** = *σ****v***, where ***v*** = (*δH, δQ*) and the matrix ℒ is

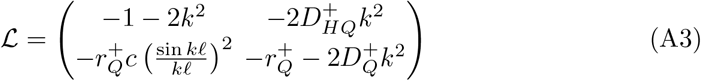

The largest growth rate is given by 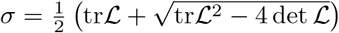, which written out explicitly is

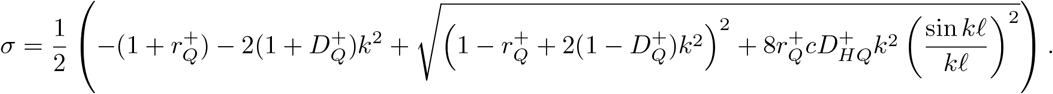

Note that because *k*ℓ *<* 1 (i.e., ℓ is smaller than the range pattern sizes), 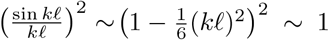. Thus for *k*ℓ *<* 1 the effects of the average ⟨*H*⟩ on the linear instability are small. This is not necessarily the case in the nonlinear regime.

If *D*_*HQ*_ or *c* vanish then *σ <* 0 always and there is no linear instability. Of course if *r*_*Q*_ = 0, nothing happens *σ* = 0. If *D*_*Q*_ = 0 there is an instability but there is no fastest growing mode because there is no stabilizing mechanism for small wavelengths (*σ* ∼ const. as *k* → ∞) (the growth rate as *k* → ∞ scales as −*D*_*Q*_*k*^2^).

We can find the modes *k*_0_ where the growth rate vanishes, *σ*(*k*_0_) = 0 (taking *k*ℓ ≪ 1):

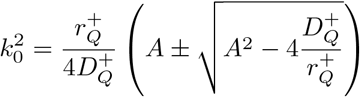

where 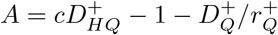.

The modes *k*_0_ bound the regime of unstable wavelengths that grow and lead to the pattern. This occurs after the bifurcation past the critical mode *k*_*c*_ where *σ* = 0 and *dσ/dk* = 0. This means that for patterns to form 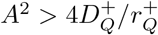 which gives

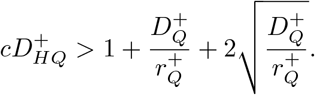

The scaling of the critical mode is found as 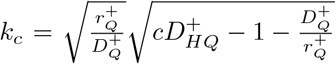. From this we find that the fastest growing wavelength *λ*^*^ (the size of the pattern) can be approximated as

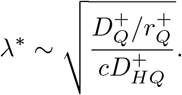

This approximation is most accurate for *k*ℓ ≪ 1 and for 0 ≲ *σ* ≲ 1 (see Fig. A1). For small increases in the length scale ℓ we expect

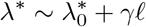

where 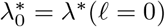 and *γ* ∼ 2.5 (Fig. A1). Note that for increasing *k*ℓ and/or 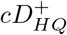 the dependence of the growth rate *σ* on ℓ increases, and thus the dependence of *λ*^*^ on ℓ becomes more complex/nonlinear.

**Fig. A1.**
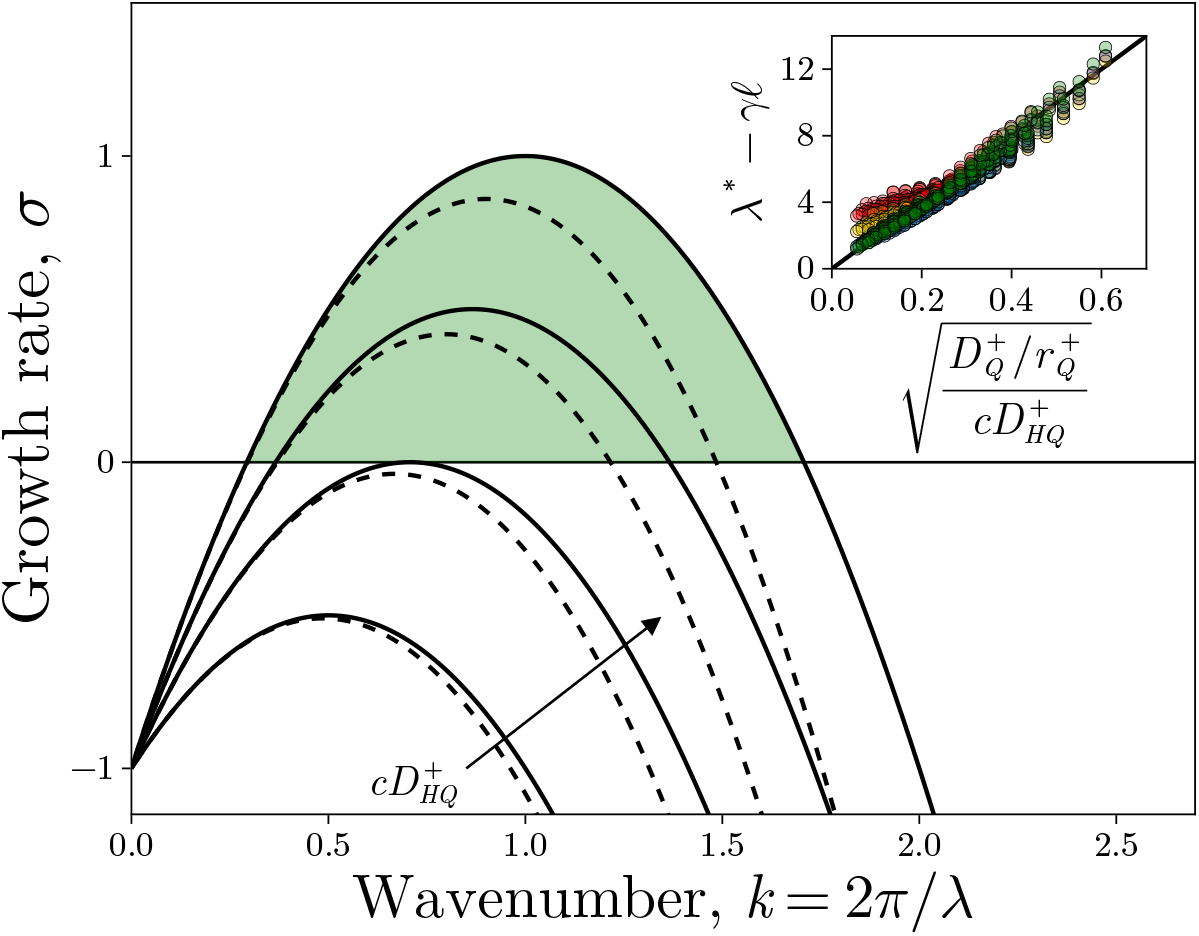
Growth rates curves *σ*(*k*) for a range of 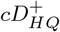 for 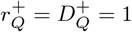 and ℓ = 0 (solid lines) and ℓ = 0.5 (dashed lines). Green region shows ranges of wavelengths where *σ >* 0 and pattern formation occurs (for ℓ = 0). Inset shows fastest growing wavelength *λ*^*^ versus the approximated scaling ratio 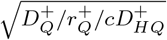 for 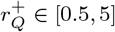 and 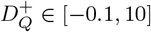. We take 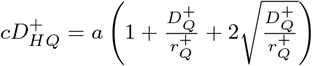 for *a* ∈ [1, 5]. *λ*^*^ is corrected by the perturbations *γ*ℓ where *γ* = 2.5 and ℓ = 0 (green), 0.25 (blue), 0.75 (gold), and 1.25 (red). Black line in the inset is a linear fit *Y* = 20*X*.

